# Intrinsic flexibility of Type VII secretion central pore is required for substrate translocation

**DOI:** 10.64898/2026.07.31.742005

**Authors:** Christina Ritter, Katherine S.H. Beckham, Grzegorz Chojnowski, Edukondalu Mullapudi, Frank Stein, Mandy Rettel, Mikhail Savitski, Matthias Wilmanns

## Abstract

Mycobacteria have up to five distinct type VII secretion pathways that play diverse roles in the cell ranging from iron uptake to virulence. To date, high-resolution structures of a hexameric ring-like pore complex have only been determined for closely related mycobacterial ESX-5 systems. The most significant difference between them is the arrangement and flexibility of the inner transmembrane helices of the EccC*_5_* subunit within the central pore, leading to either a closed or to a semi-open conformation of the pore. In this work, we probed the functional roles of several central pore-forming residues in mediating secretion and demonstrate their crucial role in ESX-5 substrate translocation efficiency. Structural characterization of an ESX-5 variant with a conserved proline (P73A) surprisingly revealed rigidification of the pore-forming helices. In summary, our data demonstrate that maintaining the conformational flexibility of the central ESX-5 pore is essential for substrate secretion. Our findings reveal how the rigid structural ESX-5 scaffold with separate sections – facing the periplasm, the inner mycobacterial cell wall membrane and facing the cytosol – provides a complex framework for a high level of pore dynamics, restricted to the inner EccC*_5_* subunit pore-forming helices. Their proper integration is required to generate a functional secretion system. The high level of conservation of several pore-forming residues across different type VII secretion systems indicates our findings to be general valid for these systems.

**SIGNIFICANCE STATEMENT:** A key virulence signature of the deadliest pathogenic mycobacteria is the presence of a number of highly specialized the type-VII secretion systems, which translocate distinct infection-promoting proteins through a central pore. Recent high-resolution structures of one of these systems (ESX-5) have revealed closed or partially closed pore states. This raises the question of how pore opening is mediated to allow substrate transport. Here, we demonstrate that this system is highly sensitive to even subtle changes within the central pore. Loss of conformational pore flexibility impairs secretion. A comprehensive understanding on the structure and associated dynamics of the central pore of type-VII secretion systems is essential to further unraveling mycobacterial virulence and developing future strategies for combatting it.

## INTRODUCTION

Bacteria rely on sophisticated secretion machineries to transport proteins across their cell envelope. These export pathways are essential for many different physiological processes, such as nutrient acquisition, adhesion to host cells, modulation of host-pathogen interactions, virulence, and viability. Mycobacteria utilize three protein secretion systems, namely the SEC system, the TAT system and the ESX/type VII secretion system (T7SS). While the SEC and TAT systems are ubiquitous, the presence of the ESX systems is species-specific. In pathogenic mycobacteria, these systems are encoded in up to five gene loci termed ESX-1 to ESX-5^1,2^. Each *esx* locus encodes the membrane-embedded structural components of the secretion machinery (EccB, EccC, EccD, EccE, MycP), cytosolic components (EccA, EspG) and some of the secreted substrates (Esx, PE/PPE proteins). T7SS-mediated secretion of Esx and PE/PPE heterodimeric complexes across the mycomembrane requires energy generated by ATP hydrolysis mediated by EccC, which belongs to the category of FtsK/SpoIIIE AAA+ ATPases^3–5^. EccC is unique amongst this ATPase superfamily as it encodes an array of three cytosolic ATPase domains at its C-terminus, which are connected to two N-terminal transmembrane helices (TMHs) via a domain of unknown function (DUF). In addition to providing energy for secretion, EccC also binds to secreted substrates^3,4,6^.

Recent cryo-electron microscopy (EM) structures of ESX systems have paved the way for achieving a molecular understanding of the secretion mechanism of this class of transport system. These structures included two dimeric subcomplexes of ESX-3 systems from *M. smegmatis* (Msmeg)^7,8^, which revealed the architecture of the protomeric subunit assemblies composed of EccC_3_, EccB_3_, EccD_3_ and EccE_3_ with a 1:1:2:1 stoichiometry. Hexameric structures of the ESX-5 system from *M. xenopi* (*Mxen*)^9^ and *M. tuberculosis* (*Mtb*)^10^ provided structural insights into the complete pore architecture, with the central secretion pore formed by EccC_5_ **(Figure 1).** The corresponding sequences are closely related and, hence, it was not surprising that major parts of the overall arrangement of the two structures are virtually identical. The main conformational differences are caused by the presence of the MycP_5_ subunit in the *Mtb* structure *versus* its absence in the *Mxen* structure. When MycP_5_ is present, a well-defined folded arrangement at the periplasmic ESX-5 segment with 2:1 EccB_5_:MycP_5_ stoichiometry and threefold symmetry is observed **(Figure 1b)**. In contrast, in the absence of MycP_5_ a more mobile V-shaped arrangement of three EccB_5_ dimers in distinct positions was found **(Figure 1a)**. In the presence of MycP_5_, the threefold symmetry of the periplasmic segment is maintained in the transmembrane pore, which consists of a trimer of [EccB_5_:EccC_5_:(EccD_5_)_2_:EccE_5_]_2_:MycP_5_ modules that leads to a closed pore arrangement **(Figure 1b, right)**^10^. In contrast, the *Mxen* ESX-5 structure without MycP_5_ exhibits six EccB_5_:EccC_5_:(EccD_5_)_2_:EccE_5_ protomers related by sixfold symmetry along the central pore axis, leading to a partially open pore arrangement with a central pore diameter of up to approximately 10 Å (**Figure 1b, left)**^9^.

**Figure 1.**
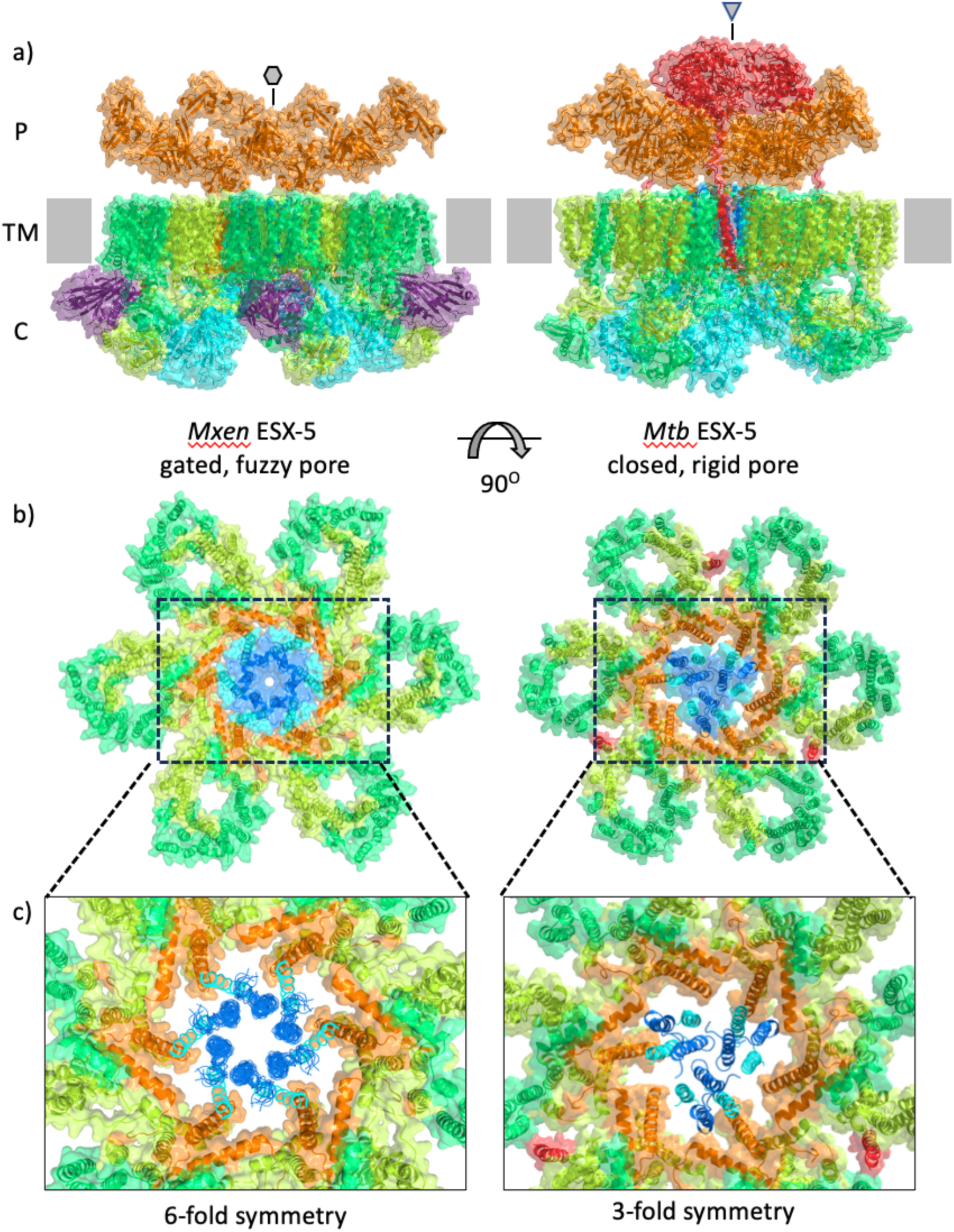
Conformational dynamics of the central ESX-5 secretion pore from *M. xenopi* (PDB ID 79BS, left) and *M. tuberculosis* (PDB ID 7NP7, right). a) Side view of the complete structural models, which include the periplasmic face section (P), the transmembrane section (TM), and the cytosolic face section (C) in combined semi-transparent surface and cartoon presentation. **b)** View from the periplasmic face along the central pore axis, demonstrating the structural conservation of the TM scaffold of different ESX-5 structures, except the central pore. For reasons of clarity, only the TM sections of EccB_5_, EccC_5_ and EccD_5_ and are shown. **c)** Zoom into the central ESX-5 pore. To illustrate the different arrangements of the EccC_5_ inner pore helices TMH1 (cyan) and TMH2 (marine blue), leading to a semi-open pore with an approximate 10 Å diameter in the *M. xenopi* ESX-5 structure and a complete pore closure in the *M. tuberculosis* ESX-5 structure, the semi-transparent surfaces of these helices have been removed. In the *M. xenopi* structure, the EccC_5_ segment is flexible and is therefore shown as an ensemble. The different symmetries along the central pore axis, observed in the two ESX-5 structures, are indicated. Color codes: EccB_5_, orange; EccC_5_, cyan (except TMH2); EccD_5_ (pore-proximal subunit), limon; EccD_5_ (pore-distal subunit), lime green; EccE_5_, violet; MycP_5_, red. Note that not all subunits or parts of them are defined in both structures.

In the *Mxen* ESX-5 structure, the central pore formed by the transmembrane helices of EccC_5_ is enclosed by the transmembrane and membrane-proximal helices of EccB_5_, referred to as the ’EccB_5_ basket’ **(Figure 2a-c**)^10^. This arrangement allows both subunits to form a strong and specific interaction site at the cytosolic TM face within each ESX-5 protomeric unit, indicating that this interaction is crucial for the overall structural organization of the holo ESX-5 pore complex. In contrast, the interface between EccB_5_ and EccC_5_ within the TM segment is formed exclusively by hydrophobic residues from the EccB_5_ TMH and the EccC_5_ TMH1 and therefore is transient in nature. Due to a tilt angle of approximately 30 degrees between these two helices, the average distance between their principal axes increases from 10 Å at the cytosolic TM face to 14 Å at the periplasmic TM face. This arrangement enables the formation of sixfold repeated outer and inner ring-like arrays by EccC_5_ TMH1 and TMH2, which constitute the ESX-5 pore **(Figure 1a, left;** **Figure 2a).** Unlike the structurally well-defined EccC_5_ TMHs that form the closed pore in the *Mtb* ESX-5 assembly with snug packing interactions between the EccC_5_ TMHs from all six protomers, the same helices are highly flexible in the *Mxen* ESX-5 structure and could only be modelled as an ensemble of multiple conformations^9^, suggesting that significant conformational dynamics are involved in the pore opening process. The data also demonstrate that the proper positioning of the EccC_5_ subunit primarily depends on its interactions mainly with the EccB_5_ subunit outside the TM segment, as opposed to transient helix/helix interactions within the TM segment **(Figure 2c).**

**Figure 2:**
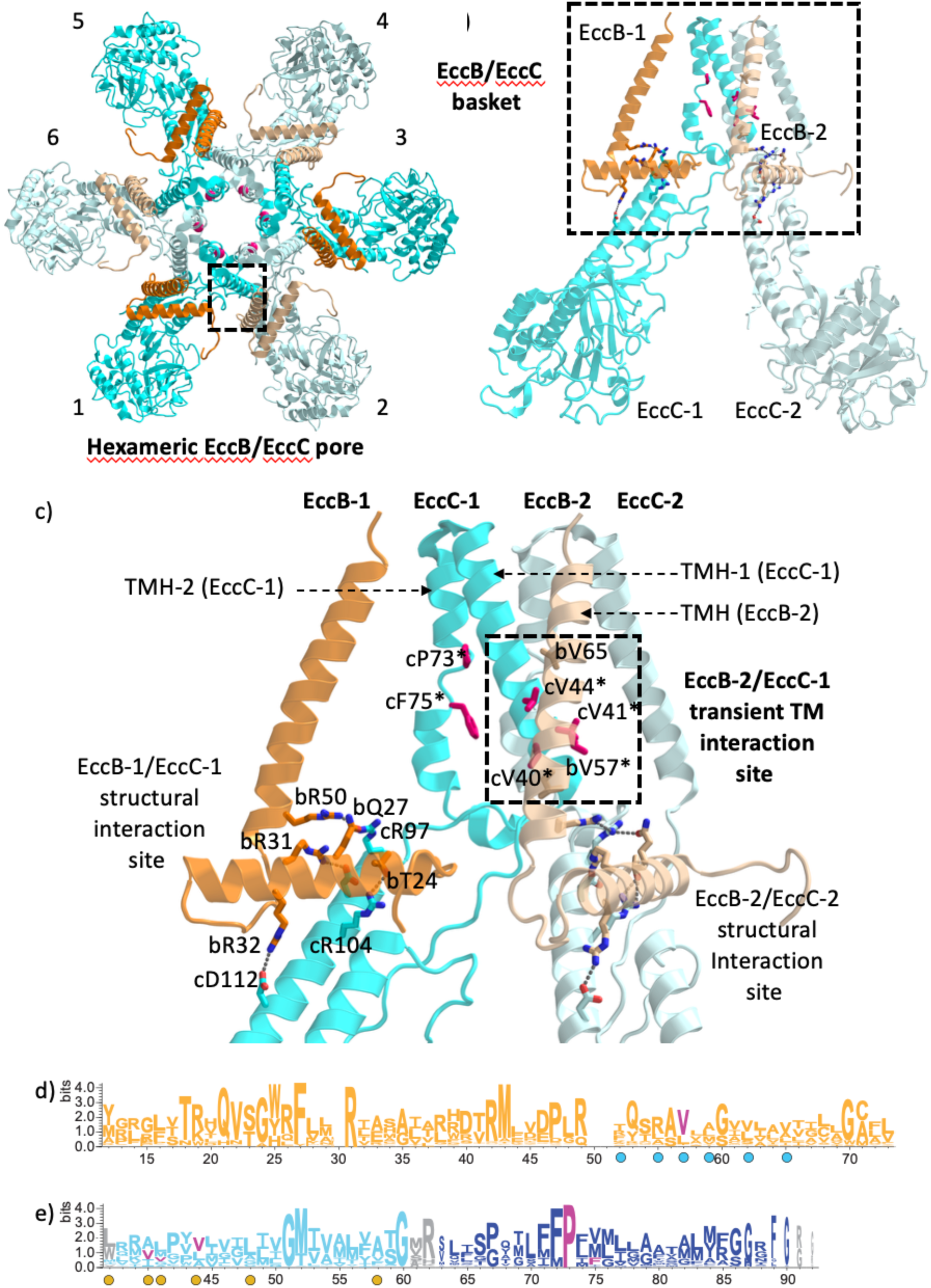
The central ESX-5 pore architecture and dynamics are established by interactions between EccB*_5_* and EccC*_5_*. **a)** Hexameric ESX-5 EccB_5_/ EccC_5_ arrangement, viewed from the cytosolic face along the central pore axis, as observed in the ESX-5 structure from *M. xenopi* (PDB ID 79BS). Neighboring EccB_5_ and EccC_5_ subunits are in alternating full/faint colors adopting conventions of Figure 1. **b)** Side view of a dimeric module extracted from the hexameric ESX-5 EccB_5_/EccC_5_ arrangement. **c)** Zoom of the side view, focusing on the bipartite cytosolic face and TM EccB_5_/ EccC_5_ interaction sites (boxed in panel b). This EccB_5_/ EccC_5_ interaction site is generated by mostly specific, polar interactions and is part of an extended interaction network also involving the two EccD_5_ subunits from the protomeric unit. This interface is highly conserved in different ESX-5 structures and is essential for the structural integrity of the overall ESX-5 scaffold^9,10^. In contrast, due to domain swapping in EccB_5,_ the second EccB_5_/ EccC_5_ interaction site within the TM segment (boxed) is generated by EccB_5_ and EccC_5_ subunits from neighboring ESX-5 protomeric units. This site is mainly of hydrophobic nature and lacks any specific interactions. Since in the *M. xenopi* ESX-5 structure, the EccC_5_ TM segment is mobile^9^, this site likely contributes to ESX-5 pore dynamics rather than to the integrity and stability of the overall structural scaffold of the ESX-5 complex. Logo presentation of relevant **d)** EccB_5_ and **e)** EccC_5_ sequence segments indicating conservation. Residues of the corresponding TM segments contributing to EccB_5_/EccC_5_ interactions are highlighted by circles in complementary colors of the interacting subunit. Color codes for EccC_5_ TMH1 and TMH2 are as in Figure 1. Residues that were mutated for functionally probing their contributions to ESX-5 pore dynamics are colored in magenta and are indicated by asterisks in c).

Based on known structures of folded substrates^11^, we proposed that a further opening of the pore to approximately 20 Å would be required and that the conformation of the *Mxen* structure, therefore, represents a partially open gated conformation^9^. The nature of the conformational changes and dynamics required to open the central pore and allow substrate secretion, remains unknown. To approach this central question, we explored the role of the EccB_5_ and EccC_5_ transmembrane helices in mediating secretion. Most ESX-5 mutants had significantly impaired for secretion efficiency. Finally, we used the ESX-5 EccC_5_ P73A (cP73A) variant, which showed only residual secretion, for structural analysis by single particle cryo-EM. Unexpectedly, we observed rigidification and straightening of the inner pore helix in this ESX-5 variant, resulting in a less dynamic pore structure than that observed in the *Mxen* wild-type (WT) ESX-5 complex. These results suggest that pore plasticity plays a fundamental role in Type VII secretion.

## RESULTS

### Experimental design of Mxen ESX-5 pore variants

In this study, we aimed to characterize structural features of the central ESX-5 TM pore, assuming that its architecture and associated dynamics are crucial for substrate secretion. As the inner ring-like structure of the ESX-5 TM pore is formed by two helices from EccC_5_ (TMH1, TMH2) and a single TM helix from EccB_5_, we probed residues from the corresponding sequence segments **(Figure 2)**. In EccC, P73 is one of the few invariant residues in all ESX systems (ESX-1 to ESX-5) **(Figure 2e)**. This proline interrupts the helical hydrogen bond pattern of the EccC_5_ TMH2 and could therefore contribute to the disorder of the inner EccC_5_ TM pore observed in the structure of the *Mxen* ESX-5 pore complex. Furthermore, the *Mxen* ESX-5 composite model ensemble shows that two phenylalanine side chains (F72, F75) in TMH2 are consistently oriented towards the inner surface of the central TM pore **(Figure 2c)**^9^. We selected P73 and F75 for site-directed mutagenesis. To examine the impact of the transient interactions between the EccB_5_ TMH and EccC_5_ TMH1 we mutated a total of four valines into phenylalanine (bV57F, cV40F, cV41F, and cV44F), which all contribute to this interface **(Figure 2c-d).**

### Mxen ESX-5 pore variants impact substrate secretion

*Mxen* ESX-5 mutant complexes were generated by modifying the pMV-ESX-5 plasmid used to produce the WT complex **(Supplementary Table S1)** and were expressed and purified using established protocols^9^. All mutant complexes were produced with similar yields and behaved as the WT complex **(Supplementary Figure S1a)**.

We then tested the impact of the mutations on the function of the ESX-5 system by monitoring the secretion of T7SS-specific substrates using a proteomics-based secretion assay. *M. smegmatis* expressing the modified pMV-ESX-5 constructs were cultured to an OD_600_ of 1 and the secreted and whole cell fractions were isolated. Using tandem mass tag labelling (TMT) coupled with quantitative mass-spectrometry we quantified the relative protein expression for each mutant, focusing on the proteins encoded on the pMV-ESX-5 construct **(Figure 3).** This design was expected to be feasible, as *M. smegmatis* does not encode an ESX-5 secretion system that could secrete the plasmid-encoded ESX-5 substrates^5,12,13^. The analysis focused on the abundance of secreted proteins present in both the culture filtrate (secretome) and in the whole cell (WC) fraction. Besides EsxM and EsxN, two PE/PPE pairs are encoded within the *Mxen* ESX-5 locus **(Figure 3a**). For simplicity, we named these proteins as PE1, PPE1, PE2 and PPE2, corresponding to the unannotated genes *mxen9349*, *mxen9354*, *mxen9364* and *mxen9369*, respectively. As part of this assay, we included two negative controls, the *M. smegmatis* MC^2^155 expression strain (WT), which does not encode the ESX-5 substrates, and the ΔEccE*_5_*/ΔEccA*_5_* mutant^5,12,14^. The ΔEccE*_5_*/ΔEccA*_5_* variant remains capable of forming a pore complex, but is unable to secrete substrates, thus serving as a negative control. These control experiments also confirm that these substrates are not secreted by any endogenous *M. smegmatis* ESX secretion system. The Liquid Chromatography-Tandem Mass Spectrometry (LC-MS/MS) data were normalized to WT ESX-5 levels.

**Figure 3.**
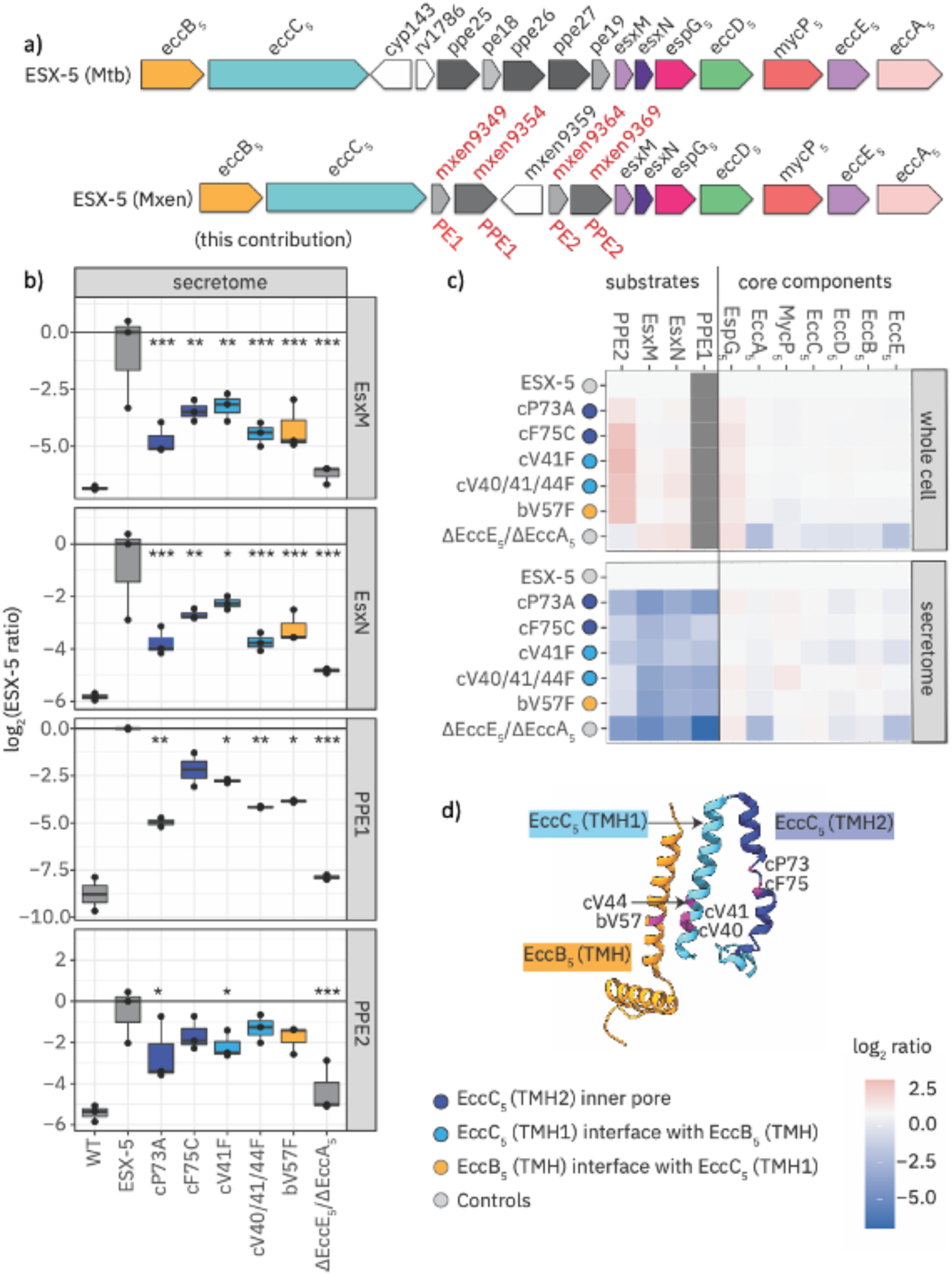
Changes in the EccC*_5_* TM section and the EccB*_5/_*EccC*_5_* TM interface impair ESX-5 secretion. **a)** ESX-5 operon arrangement in *M. xenopi* and *M. tuberculosis* with related ppe/pe and esxM/esxN genes coding for heterodimeric coiled coil substrates, however in different orders. To simplify referencing the relevant literature, we renamed genes mxen9349/mxen9354 and mxen9364/mxen9369 to PE1/PPE1 and PE2/PPE2, respectively. **b)** The protein abundance of four secreted substrates (EsxM, EsxN, PPE1 and PPE2) in the secreted fraction is shown for the different ESX-5 variants, normalized to ESX-5 levels. Color codes are as in Figure 1, controls are in grey. The horizontal black line represents the mean level of secretion from the WT construct (pMV-ESX-5). Statistical analysis of each variant with three independent measurements (N = 3) was performed with limma using Bayes moderated t-tests with significance levels shown: *, p < 0.06; **, p < 0.01; *** p, < 0.001. **c)** Heat map showing the relative levels of all the plasmid encoded *M. xenopi* ESX-5 components, calculated based on log_2_ ratio change in protein levels of all respective mutant complexes compared to the pMV-ESX-5 ‘WT’ control. Negative-fold changes (blue) indicate a decrease in protein levels in either the whole cell fraction or the culture filtrate (secretome); positive fold changes are shown in red. Where the protein was not detected due to low abundance, a grey box has been shown**. d)** Overview of mutants mapped on EccB_5_/EccC_5_ TMD regions (*cf.* Figure 2c).

Our data revealed a significant decrease in the secretion of EsxM, EsxN, PPE1 and PPE2 in all ESX-5 mutants **(Figure 3b)**. On the other hand, the levels of PE1 and PE2 were too low for quantitative detection in this assay and were therefore excluded from further analysis. For all substrates tested, secretion was most diminished in the cP73A ESX-5 variant. In contrast, the levels of the structural components remained unchanged in the WC fraction, indicating that these point mutations did not disrupt structural integrity of ESX-5 complex formation **(Figure 3c).** Only in the ΔEccE*_5_*/ΔEccA*_5_* ESX-5 variant, we observed an overall decrease in the levels of structural ESX-5 subunits, indicating that the structural integrity was compromised. We also observed a slight increase in the levels of the tested substrates and the cytosolic PE/PPE chaperone EspG in the WC fraction of most ESX-5 variants, indicating that they may accumulate in the cytosol due to reduced secretion. The level of PPE1 in the whole cell fraction was insufficient for quantification. Overall, all of the mutants designed based on our *Mxen* ESX-5 pore complex structure exhibited significant defects in ESX-5 substrate secretion. As the cP73A ESX-5 mutant showed the strongest effect on secretion deficiency we undertook further structural characterization of this ESX-5 variant.

### Structural analysis of the cP73A ESX-5 variant reveals pore rigidification

To obtain a sample suitable for structural analysis, the cP73A ESX-5 mutant complex was further concentrated by gradient centrifugation to obtain sufficiently pure hexameric complex. Blue Native (BN)-PAGE and Negative stain (NS)-EM revealed a large monodisperse species as observed for the WT ESX-5 complex^9^ (**Figure S1b-c).**

Using single particle cryo-EM, a total of 10,233 movies were collected and subsequently processed both without applying symmetry (C1) and with imposed C6 symmetry, assuming that the sixfold symmetry observed in the *Mxen* WT ESX-5 structure was maintained^9^. While the resolution of the C1 map was limited to 5.3 Å, the C6 map reached a global resolution of 3.9 Å **(Figure 4a-b, Figure S2**, **Supplementary Table S2).**

**Figure 4:**
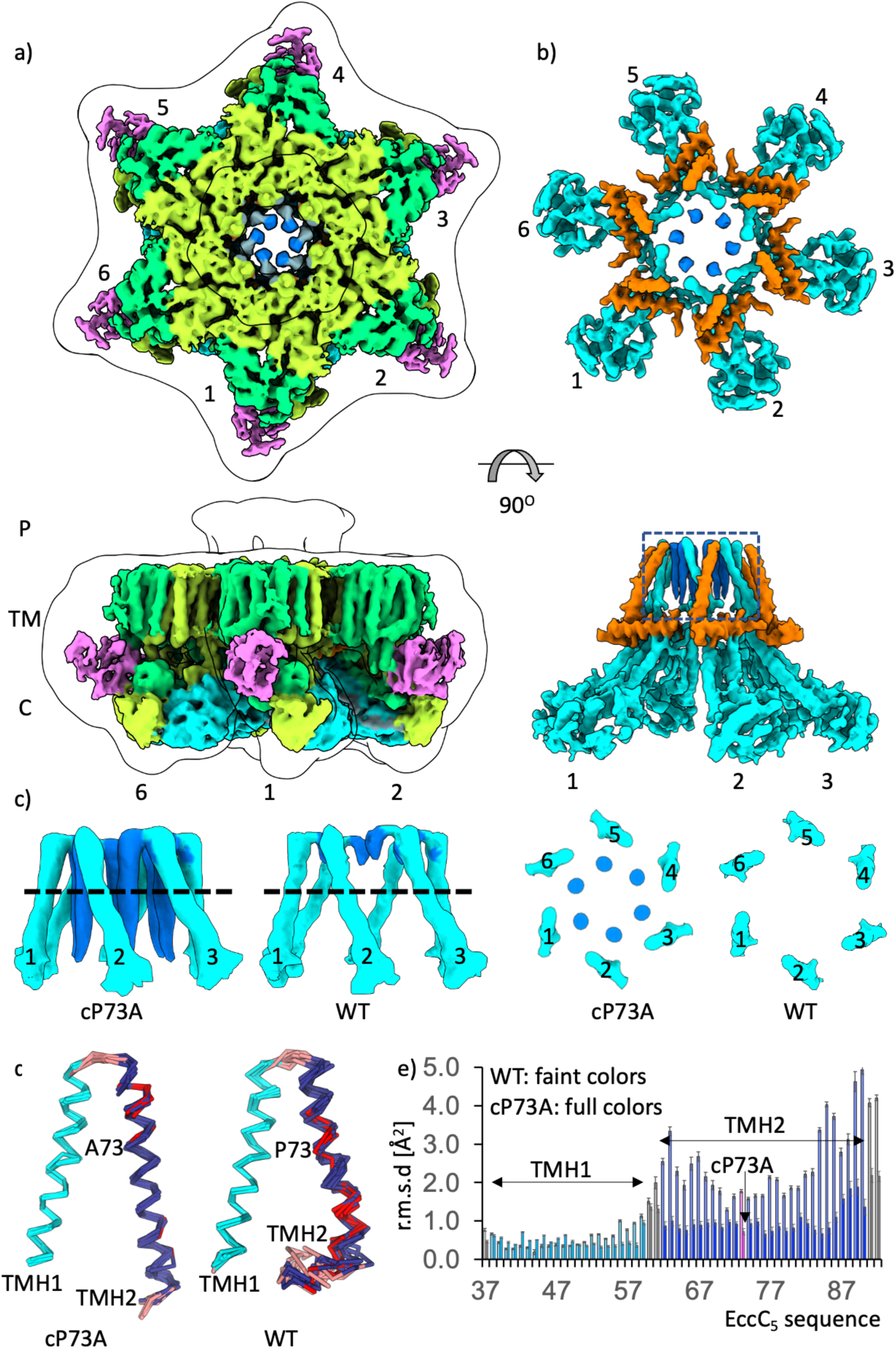
Changes in the ESX-5 pore architecture and dynamics induced by the cP73A mutation. Electron microscopy maps of **a)** the cP73A ESX-5 complex and **b)** the same complex restricted to EccB_5_ and EccC_5_ in top and side view. Note that EccB_5_ and EccC_5_ are mostly hidden by the surfaces of other ESX-5 subunits in the top view. An additional contour line (contour level 0.13) indicates the boundaries of a low-pass filtered map. Color codes are as in Figure 1. The approximate positions of the sixfold-repeated ESX-5 protomeric EccB_5_/EccC_5_/EccD_5_(proximal)/EccD_5_(distal)/ EccE_5_ units are numbered when visible. **c)** Zoom of the density accounting for the cP73A ESX-5 EccC_5_ TM segment (boxed in panel b) in top and side view. For comparison, the density accounting for the WT ESX-5 EccC_5_ TM segment is also shown. To highlight the lack of visibility of most of the WT EccC_5_ TMH2, the side views are sliced, as indicated by a dashed line in the side view. The approximate positions of the sixfold repeated EccC_5_ TMH1s are numbered when visible. The inner pore ring is formed by the sixfold repeated EccC_5_ TMH2s (marine blue) to an extent they are visible. **d)** Superposition of the top 10 Rosetta models of the cP73A and WT EccC_5_ TM segments, indicating reduced conformational diversity especially of the EccC_5_ TMH2 in the cP73A ESX-5 structure. Segments lacking helical conformation are shown in red, when located within TMH1 and TMH2 segments, and in salmon when located at the terminal positions of the TM segment. The positions of EccC_5_ P73 and A73 (cP73A) are indicated. e) R.m.s.d. values of the EccC_5_ TM segment of WT and cP73A ESX-5. Error bars depict standard errors of the mean from a bootstrap analysis.

**Figure 5.**
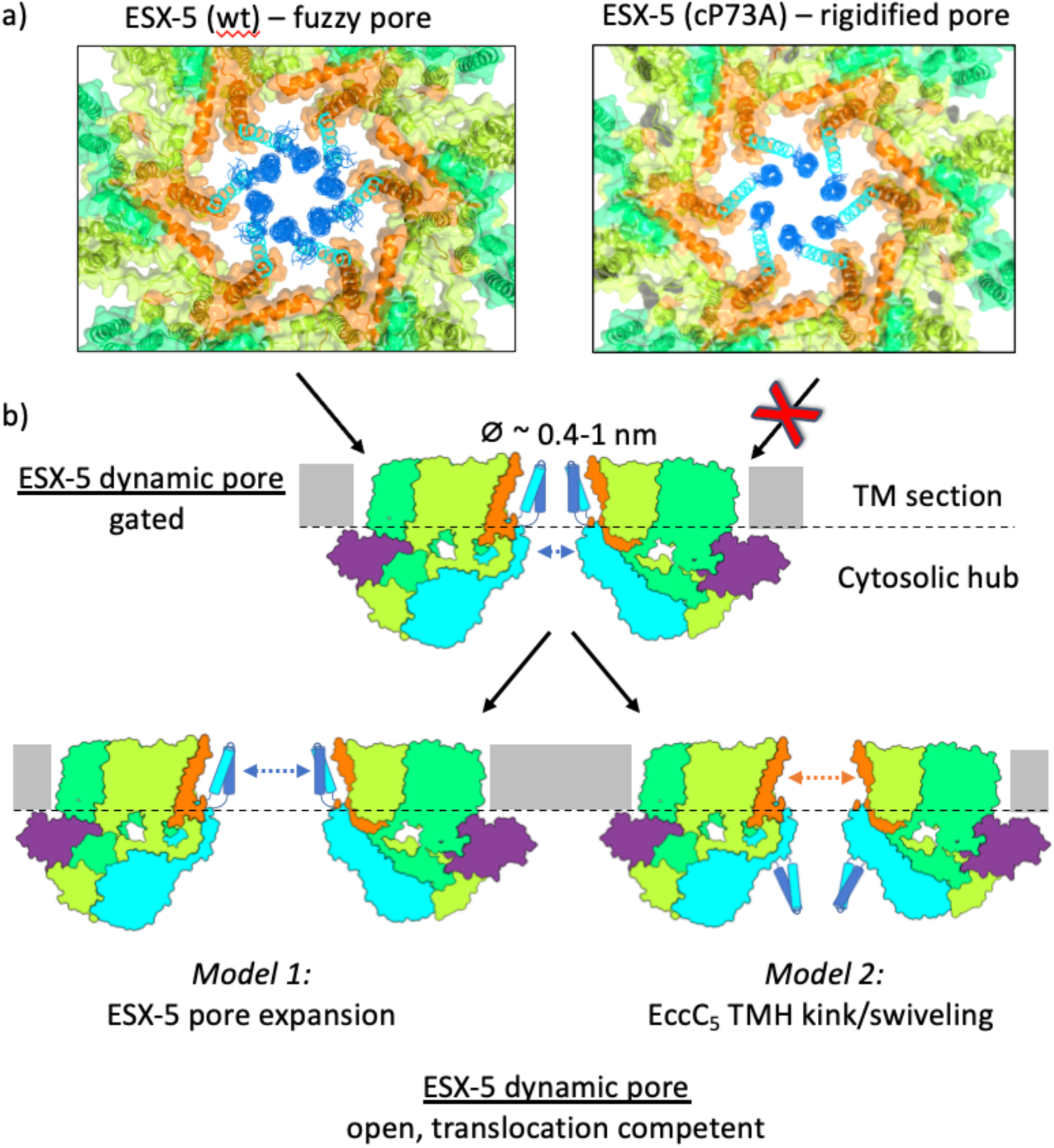
Models of pore expansion, allowing substrate secretion. **a)** Structural comparison of the central pore of the WT and cP73A ESX-5, using the layout and color codes of Figure 1. Rigidification of the EccC_5_ TM section in cP73A ESX-5 structure (right) impairs the dynamics of the fuzzy pore, observed in the WT ESX-5 structure (left). Note that the observed gated pore conformation in the WT ESX-5 structure does not have a sufficient diameter to allow translocation of folded ESX-5 substrates. Insight into the type and mechanism of any required conformational changes for further pore opening are currently still unknown. Our data indicate that impairment of EccC5-mediated pore dynamics most will likely also block any further conformational changes for pore opening. **b)** Hypothetical schematic models on ESX-5 pore opening, leading to a translocation-competent ESX-5 pore conformation. ESX-5 pore expansion without major conformational changes of the EccC_5_ TM section (model 1) would require disrupting specific interactions between the ESX-5 structural scaffold components EccB_5_, EccC_5_ and EccD_5_ (*cf*. Figure 2c). In contrast, opening of the ESX-5 pore conformational changes in the EccC_5_ TM section, here proposed as TM kinking/swiveling (model 2), would not necessarily affect specific interactions between the ESX-5 structural scaffold components EccB_5_, EccC_5_ and EccD_5_. The purpose of these models is to inspire future research questions.

The cP73A ESX-5 complex displayed the same overall structural architecture as the WT ESX-5 complex **(Figure 4a)**. In particular, the inner EccC_5_ pore helices were arranged similarly, with the EccC_5_ TMH1 forming the outer layer of the pore through interactions with the EccB_5_ TMH, generating the established EccB_5_/EccC_5_ basket arrangement, and TMH2 forming the inner face of the central pore^9^ **(Figure 4b)**. However, unlike the WT ESX-5 pore complex, the density of TMH2 in the cP73A ESX-5 mutant was significantly enhanced, revealing structural details of this helix not observed in the WT ESX-5 structure (**Figure 4c)**. Only in the cP73A ESX-5 structure, the sixfold repeated EccC_5_ TMH2s appear as vertical stalactite-like densities in the side view of the overall structure with connections to the sixfold repeated TMH1’s at the periplasmic face. In contrast, in the WT ESX-5 structure weak density only becomes visible by low-pass filtering. In a top view of the cP73A ESX-5 EccC_5_ TM section, sliced about halfway through, the difference in TMH2 visibility between the cP73A and WT ESX-5 structures is most apparent **(Figure 4c)**. These differences were further supported by a difference map filtered to 5 Å resolution^15,16^ and a local cross-correlation map from half-maps of the cP73A and WT ESX-5 reconstructions (**Figure S3)**.

To further probe the structural impact of the cP73A mutation on the conformational dynamics of the TMH2, we generated an ensemble of models to fit into the TMH2 density, using a Rosetta-based approach previously used for the WT ESX-5 complex^9^. By analyzing 100 ensemble structures, we observed an overall reduction in residue root mean squares deviation (r.m.s.d.) values of the entire EccC_5_ inner pore segment from 2.06 Å (WT) to 0.97 Å (cP73A) **(Figure 4d-e).** These differences are most pronounced in the second half of the EccC_5_ TM segment covering the inner pore TMH2. From these findings, we conclude that P73 plays a pivotal role in driving conformational flexibility in the ESX-5 pore, which is essential for protein secretion.

## DISCUSSION

High-resolution structures of mycobacterial T7SS pore complexes are currently limited to the ESX-3 and ESX-5 systems. Although major principles of the underlying architecture are preserved in these structures, they also reveal astonishing differences that cannot be explained by sequence and functional diversity alone. One major difference is in the conformational diversity and associated flexibility of the TM segment of EccC_5_. Whereas in the dimeric ESX-3 structures, this segment is invisible likely due to its flexibility, in the two known hexameric ESX-5 complexes the same segment is either partially flexible or rigidified, depending on the conformation it adopts within the central pore **(Figure 1)**. Taken together, these data suggest that the EccC_5_ TM section presents a mobile element that is not part of the scaffold of the remaining TM ESX-5 subunits (mainly EccB_5_ and EccD_5_), comprising rigid and well-defined TM helices. This is supported by our observations that all specific EccC_5_/EccB_5_ interactions are confined to sequence segments outside the TM section (**Figure 2c).** These data also explain why, despite its loose interactions within the TM segment, EccC_5_ is essential for the overall structural assembly of ESX pore complexes^5,13^. Since any transient EccB_5/_EccC_5_ TM interactions are only observed within pore-forming hexameric ESX assemblies, our data demonstrate that the assembly competent for secretion is hexameric, which is a common feature of FtsK-AAA+ systems^3,17^. Known T7SS substrates share a common structural motif consisting of an antiparallel four-helical bundle. However, the sequence of these proteins can range from 100 to 1400 residues in length^18^, suggesting that the transport system requires an additional degree of flexibility to accommodate such variation. Previous structural data have shown that the cytoplasmic domains of EccC_5_ display a high degree of conformational flexibility, mediated by loop segments between the domain of unknown function (DUF) and threefold repeated arrays of ATPase domains^3,12^. Here, we have shown that ESX-5 structure and function are also highly sensitive to EccC_5_ TMH dynamics. Based on our assumption that a TM pore size of at least 20 Å would be required to allow ESX-mediated secretion of folded substrate^9^, several questions arise about the molecular mechanisms required to achieve a level of pore opening permissive for ESX-5 substrate translocation.

One plausible model for pore opening could arise from the intrinsic flexibility observed in the complete ESX architecture, which has been shown to be interchangeable between hexameric TM arrangements with either sixfold symmetry, leading to equal interactions between neighboring protomeric assemblies, and threefold symmetry, inducing an alternating pattern of uneven interactions between neighboring protomeric assemblies **(Figure 1).** The latter arrangement results in a trimer of dimers, which has also been observed in separate ESX-3 dimers^7^. In addition to the EccC_5_ subunit, in particular the TM segments of EccE_5_ and MycP_5_ exhibit varying degrees of mobility within these structures or are even absent, unlike EccB_5_ and the two copies of EccD_5_ that have well-defined TM segment arrangements in all available structures. Alternatively, one may consider are kink/swivel motions of the EccC_5_ TMHs within or towards the outside of the TM section, originating from the EccC_5_ stalk that is embedded in a large network of specific interactions, especially with the N-terminal segment of EccB_5_ **(Figure 2c)**. Related motions have been shown to play a key role in the activity of several transmembrane proteins^19,20^. The work presented here suggests that this flexibility is required for the efficient secretion of protein substrates by the T7SS. Based on the presented data, it is plausible to conclude that rigidification of the EccC_5_ TM segment reduces the ability of the EccB_5_ TMH to adopt variable arrangements with other interacting ESX-5 TM subunits, impacting the overall TM segment plasticity in these systems.

As the currently known protomeric unit scaffolds of the different ESX systems are structurally conserved as are the sequences of the inner pore TM segments of EccB and EccC **(Figure 2d-e),** it is reasonable to assume that our findings are applicable to all five ESX-5 secretion systems. Considering the recently published structural data, our work is a first step towards understanding the molecular mechanics by which the T7SS secretion system mediates substrate transport across the inner membrane.

## METHODS

### Cloning of ESX-5 mutants

The pMV-WT ESX-5 and ΔEccE/ΔEccA ESX-5 complexes were constructed as previously described^12^. The mutants used in this study (**Supplementary Table S1)** were created by excising fragments of the EccB/EccC_5_ genes from pMV-ESX-5 using the PacI/AfiII restriction sites and inserting them into the pJet vector (Thermo Fisher Scientific) for a minimal vector backbone. Mutants in EccB_5_ and EccC_5_ were generated by site-directed mutagenesis. All mutant sequences were verified before subcloning the EccB/EccC_5_ fragment back into the pMV-ESX-5 plasmid. The final plasmid sequences were again verified to ensure that no mutations were introduced following ligation.

### Sequence Analyses

Mycobacterial EccB and EccC sequences were obtained from the Uniprot database^22^ by selecting hits with > 50 % sequence identity to the proteins found in *Mtb (H37Rv)*. This approach retrieved between 100-200 sequences for EccB and EccC from each of the ESX systems (ESX-1 to ESX-5). Sequences were aligned using SnapGene software (SnapGene) and the Clustal algorithm^21^. Regions of the alignments were uploaded to the WebLogo server^22^.

### Expression and Purification of ESX-5 complexes

The ESX-5 and the mutant complexes were expressed and purified as previously described^9^. For cryo-EM, the cP73A complex was dialyzed against 20 mM Tris (pH 8.5) and 150 mM NaCl for two hours using a 3.5 kDa MWCO dialysis cassette (Roth) to remove any remaining detergent. The sample was then applied on top of a linear 10%-40% glycerol gradient and centrifuged at 25,000 rpm for 17.5 hours. The gradient was fractionated and screened for the presence of hexameric complexes using Blue Native-PAGE (Life Technologies) and NS-EM.

### Secretion assay

For MS based secretion assays, the constructs were transformed into *M. smegmatis* mc^2^155 groEL1 ΔC^14^. For each construct, four colonies were picked and precultures were set up. Then, 0.5 ml of the saturated precultures were used to inoculate 50 ml 7H9 medium. Cells were grown to an OD of 1 and pelleted by centrifugation at 4,000 rpm for 30 minutes. For the whole cell fraction, pellets were dissolved in PBS supplemented with 0.01 mg/ml DNAse and incubated on a rotating wheel for one hour. Afterwards, the sample was lysed using a sonicator. To ensure sufficient cell lysis and to keep membrane proteins soluble Triton X-100 was added to a final concentration of 1%, and SDS was added to a concentration of 0.1%. The suspension was then incubated for one hour. The concentration of the supernatant was adjusted to 0.4 mg/ml by a Bradford assay (Roth). For the secretome fraction, the culture supernatant was filtered through a 0.2-μm filter and the secreted proteins were precipitated by adding trichloroacetic acid (Roth) to a final concentration of 5%. Precipitated protein was pelleted by centrifugation at 7,000 rpm and 4 *^◦^*C, and then washed with acetone. The sample was then pelleted at 13,000 rpm. The supernatant was discarded and the sample air dried. The pellet was resuspended in 1 M Tris and 0.1% SDS (Sigma). For MS analysis the sample was adjusted to a concentration of 0.4 mg/ml using a Bradford assay (Roth).

### Mass spectrometry

Robust quantification of the sample was achieved by MS through isobaric labelling with Tandem Mass Tags (TMT). This labelling method allowed for the simultaneous measurement of all secreted fractions and whole cell fractions in a single LC-MS/MS run^23^. Samples were prepared using the Single-Pot Solid-Phase-Enhanced Sample Preparation (SP3) protocol^24,25^ and subjected to overnight trypsin digestion. Digested peptides were isolated and labelled with TMT10plex Isobaric Label Reagent (ThermoFisher) according to the manufacturer’s instructions. Samples were combined for TMT10plex (ThermoFisher) labelling, followed by cleanup up the sample using an OASIS® HLB μElution Plate (Waters). Offline high-pH reverse-phase fractionation was carried out on an 1200 Infinity high-performance liquid chromatography system (Agilent), equipped with a Gemini C18 column (3 μm, 110 Å, 100 x 1.0 mm, Phenomenex)^26^.

An UltiMate 3000 RSLC nano LC system (Dionex) fitted with a trapping cartridge (µ-Precolumn C18 PepMap 100, 5µm, 300 µm ID x 5 mm, 100 Å) and an analytical nanoEase™ M/Z HSS T3 C18 column 75 µm x 250 mm, 1.8 µm, 100 Å column (Waters) was coupled to an Orbitrap Fusion Lumos (ThermoFisher) via a 360 μm OD x 20 μm ID; 10 μm tip Pico-Tip Emitter (CoAnn Technologies). An applied spray voltage of 2.4 kV was used with the capillary temperature set at 275°C. A full mass scan was acquired in profile mode in the Orbitrap with a mass range of 375-1500 m/z and a resolution of 120,000. The maximum filling time was set to 50 ms with a limitation of 40,000 ions. Data-dependent acquisition was performed with the Orbitrap resolution set to 30,000, with a fill time of 94 ms and a limitation of 10,000 ions. A normalized collision energy of 38 % was applied. MS^2^ data were acquired in profile mode.

The acquired data were processed using IsobarQuant^27^ and Mascot v2.2.07 (Matrix Science), then searched against an UNIPROT Msmeg (UP000000757) proteome database that included expressed sequences, common contaminants and reversed sequences. The following modifications were included in the search parameters: Carbamidomethyl and TMT10plex (K) (ThermoFisher) were set as fixed modifications, and acetyl (protein N-terminus), oxidation (M) and TMT10plex (N-term) (ThermoFisher) were set as variable modifications. A mass error tolerance of 10 ppm was set for the full scans (MS1), and a mass error tolerance of 0.02 Da was set for MS/MS (MS^2^) spectra. Further parameters were set: Trypsin was selected as protease with a maximum of two missed cleavages allowed; a minimum peptide length of seven amino acids was required, and at least two unique peptides were required for a protein identification. The false discovery rate on peptide and protein levels was set to 0.01.

For the statistical analysis of the secretion assay, raw IsobarQuant output files were processed using R scripts. Contaminants were filtered out, and only proteins quantified with at least two unique peptides were considered in the analysis. log_2_-transformed TMT10plex reporter ion intensities (‘signal_sum’ columns) were first cleaned for batch effects using the ‘removeBatchEffects’ function of the limma package^28^ and then normalized using variance stabilization^29^. Proteins were tested for differential expression using the limma package. Replicate information was added as a factor to the design matrix given as an argument to the ‘lmFit’ function of limma. Fold changes were calculated on a log_2_ scale after normalizing protein abundance to WT ESX-5 values by calculating the difference of the mean between mutant cP73A ESX-5 and WT ESX-5 measurements.

### Negative stain-EM, Cryo-EM and cryo-EM data processing

For Negative Stain (NS)-EM, 4 μl of the sample was applied on to a 300-mesh copper grid, that was manually coated with a continuous 10 nm layer of carbon. Samples were washed twice with water and once with a 2% aqueous uranyl acetate solution that was also used in the final staining step. For cryo-EM analysis of the *Mxen* cP73A ESX-5 mutant, 4 μl of the sample was applied to a freshly glow-discharged R1.2/1.3 Cu 200 mesh grid (Quantifoil). The sample was vitrified using a GP2 plunge freezer (Leica) at 99 % humidity and 10° C, and the liquid was blotted away for 2 s before being plunged into a liquid propane/ ethane mixture.

Data on the cP73A ESX-5 mutant were collected on a Titan Krios microscope (ThermoFischer) equipped, with a K3 direct electron detector (Gatan) and a BioQuantum K3 energy filter (Gatan) at an acceleration voltage of 300 kV. Data acquisition was performed using the EPU program (ThermoFischer). A total of 10,233 movies were collected with 40 frames and a total dose of 40.92 e^-^/Å^2^ in electron counting mode. The pixel size of the movies was set to 0.85 Å.

Motion correction and Contrast Transfer Function (CTF) estimation and all further processing was performed using CryoSPARC^30.^ Using CTF-fit parameters and local-motion distances as quality criteria, 9,275 micrographs were selected from 10,233 initial micrographs for further processing. Initial templates for template-based particle picking were generated by applying the blob picker and 2D classes were generated from the extracted particles. After inspecting the selected particles using template-based particle picking, 985,743 particles were extracted from the micrographs and binned twice. Several rounds of 2D classifications were then performed. As with other datasets, the 2D classes were curated after each round of classification to ensure that only classes corresponding to intact protein were included for further processing. An *ab-initio* reconstruction was performed with the remaining 175,048 particles, requesting two classes. Only one of these classes yielding hexameric features were used for another 2D classification. The 93,495 best particles were then used for another *ab-initio* reconstruction, resulting in an unambiguous hexameric reconstruction. The particles were re-extracted without binning, and further rounds of 2D classification were performed, leading to a final set of 78,699 particles that were used to generate a new *ab initio* model. This model was used for non-uniform refinement, applying C1 or C6 symmetry restraints.

### cP73A ESX-5 map analysis

Global difference maps of cP73A and WT ESX-5 were calculated using the difference map tool implemented in CCPEM^15^, after filtering the unsharpened maps of cP73A and WT ESX-5 to 5 Å. The cutoff for fractional differences was set to 0.2 and a dust filter, and a dust size probability of 0.2 was applied. Local correlation coefficients corresponding to local signal-to-noise ratios were calculated for the WT and cP73A half-maps using EMDA^31^ with default parameters.

### cP73A ESX-5 model building

The model was built starting from an earlier cryo-EM structure of the WT complex (PDB ID: 7b9s) as a template. First, a cP73A ESX-5 protomer model was predicted using AlphaFold2^32^ with the WT structure as a template using gapTrick with default parameters^33^. This improved the local geometry of the WT model and avoided propagating errors to the new model. To remove any bias in the mutated region, the complete EccC_5_ TMH2 was removed from the WT template. This procedure resulted in a prediction with high overall confidence scores (pTM=0.81, pLDDT=83) that was subsequently rigid-body fitted to the map and refined in real space with self-restraints generated at 5Å cut-off in COOT^34^. Any local map-fit and geometry issues in the model were resolved in ISOLDE^35^. The resulting cP73A ESX-5 protomer model was refined using CCPEM^15^ and Servalcat^36^ with jelly-body restraints and C6 point-group constraints against unsharpened half-maps. Model statistics are shown **Supplementary Table 1**.

### Structural analysis of ESX-5 pore

To map the conformational space of the cP73A ESX-5 TMH2, an ensemble of 100 models was generated using density-guided enumerative sampling refinement with C6 symmetry restraints implemented in Rosetta^37^. Root-mean-square deviation was calculated for the corresponding CA atoms relative to the coordinates of the top-scoring model. HELAN^38^ was used to analyze structural irregularities in the WT and cP73A ESX-5 structures.

## DATA AVAILABILITY

The proteomics data presented in this study are available via ProteomeXchange (www.proteomexchange.org) with identifier PXD048552. The cryo-EM density maps used in this study are available under accession codes EMD-59053. The corresponding atomic model coordinates are accessible under the PDB ID 32PF, (Extended PDB ID pdb_000032PF).

## AUTHOR CONTRIBUTIONS

C.R, K.S.H.B. and M.W. designed the project. C.R., K.S.H.B., E.M. and M.R. performed the experimental work. C.R., K.S.H.B., G,C., E.M., F.S. and M.W. analyzed the data. C.R., K.S.H.B. and M.W. wrote the paper. K.S.H.B., M.S. and M.W. supported the project.

## COMPETING INTERESTS

The authors declare no competing interests.

## ACKNOWLEDGEMENTS

We acknowledge access to the EM facility at the Centre for Structure Systems Biology, Hamburg, for sample preparation, screening and data collection. We acknowledge the Proteomics Core Facility of the European Molecular Biology Laboratory (EMBL), Heidelberg, and the technical support provided by the Sample Preparation and Characterization facility at the EMBL, Hamburg. We thank Jan Rainey from Dalhousie University, Halifax, Canada, for access to the MC-HELAN server. K.S.H.B. was supported by the Royal Society [grant number URF-R1-231106].

## ABBREVIATIONS

Type VII secretion system, T7SS; *M. smegmatis*, *Msmeg*; *M. xenopi*, *Mxen*; *M. tuberculosis, Mtb*; wild type, WT; whole cell, WC; Transmembrane helices, TMHs; EccC_5_ transmembrane helix 1, TMH1; EccC_5_ transmembrane helix 2, TMH2,

## Notes

### Competing Interest Statement

The authors have declared no competing interest.

