## Supplemental Data for "Intrinsic flexibility of Type VII secretion central pore is required for substrate translocation"

### SUPPLEMENT

### SUPPLEMENTARY TABLES

**Supplementary Table 1. Plasmids used in this study.**

| Name | Description | Reference |
|---|---|---|
| pMV-ESX-5 | Encodes *M. xenopi* ESX-5 WT locus | [1] |
| ΔEccE/ΔEccA | Deletion of *eccE* and e*ccA* from of pMV-ESX-5 | [1] |
| cP73A | Point mutation in pMV-ESX in $EccC_5$ : proline 73 to alanine | This study |
| cF75C | Point mutation in pMV-ESX in $EccC_5$: phenylalanine 73 to cysteine | This study |
| cV41F | Point mutation in pMV-ESX in $EccC_5$: valine 41 to phenylalanine | This study |
| bV57F | Point mutation in pMV-ESX in $EccB_5$: valine 57 to phenylalanine | This study |
| c(V40F, V41F, V44F) | Triple point mutation in pMV-ESX in $EccC_5$: valine 40 to phenylalanine, valine 41 to phenylalanine, valine 44 to phenylalanine | This study |

**Supplementary Table 2. Cryo-EM data collection and model building statistics.**

| | [EMD-59053] [PDB ID 32PF] |
|---|---|
| **Data collection and processing** | |
| Magnification | 105,000 |
| Voltage (kV) | 300 |
| Electron exposure (e–/Å$^2$) | 40.9 |
| Pixel size (Å) | 0.85 |
| Symmetry imposed | C6 |
| Initial particle images (no.) | 985,743 |
| Final particle images (no.) | 78,699 |
| Map resolution (Å) | 3.9 |
| FSC threshold | 0.143 |
| Map resolution range (Å) | 2.5–44.6 |
| **Refinement** | |
| Initial model used (PDB code) | 7B9F |
| Map sharpening B factor (Å$^2$) | 0 |
| Map CC (mask) | 0.84 |
| **Model composition** | |
| Non-hydrogen atoms | 12,675 |
| Protein residues | 1,660 |
| Ligands | 0 |
| **B factors (Å$^2$)** | |
| Protein | 270 |
| **R.m.s. deviations** | |
| Bond lengths (Å) | 0.01 |
| Bond angles (°) | 1.62 |

| | |
|---|---|
| **Validation** | |
| MolProbity score | 1.34 |
| Clashscore | 2.66 |
| Poor rotamers (%) | 0.75 |
| **Ramachandran plot** | |
| Favored (%) | 96 |
| Allowed (%) | 4 |
| Disallowed (%) | 0 |

**SUPPLEMENTARY FIGURES**

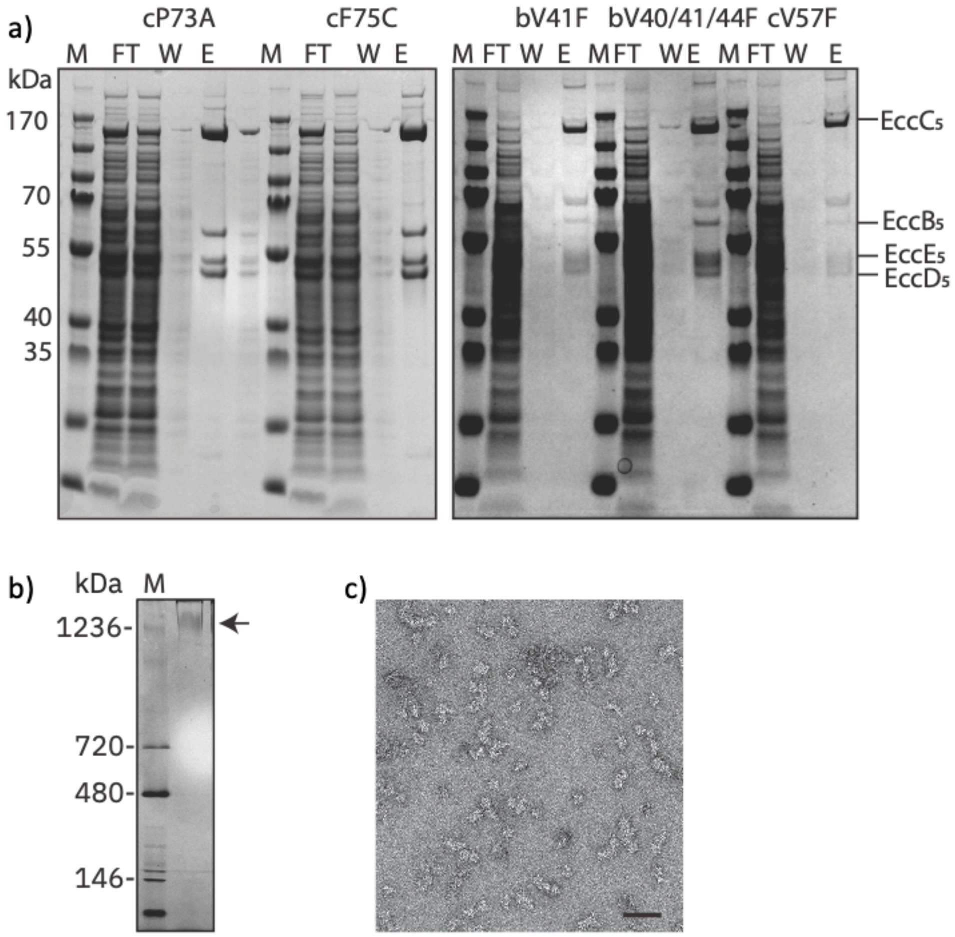


**Supplementary Figure S1 (supporting Figure 3)**. $EccB_5$ and $EccC_5$ mutations do not affect the integrity of the ESX-5 assembly. **a)** SDS-PAGE of fractions following Strep-tag affinity purification of ESX-5 variants (FT, flow-through; W, wash; E, elution). The bands corresponding to specific ESX-5 subunits are indicated on the right. **b)** Blue-Native gel showing purified cP73A ESX-5 complex. **c)** Negative stain EM image of cP73A ESX-5 complex showing the presence of ~25 nm particles, confirming the structural integrity of the ESX-5 assembly. Bar = 100 nm.

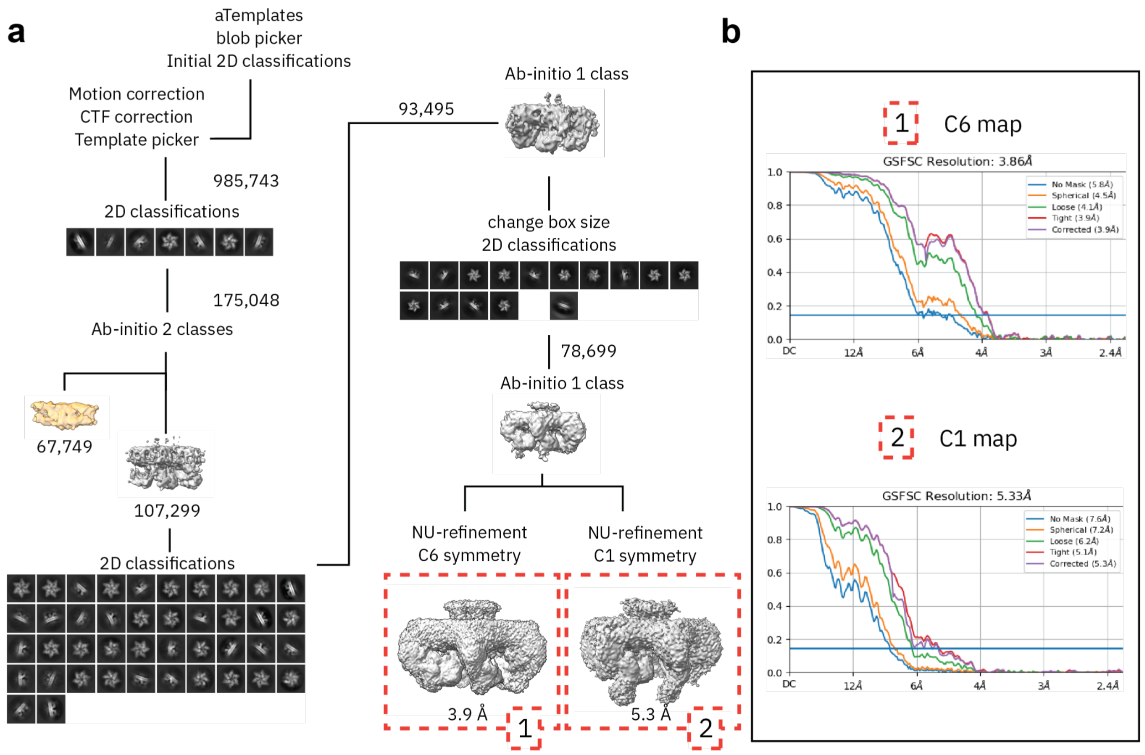


**Supplementary Figure 2**. **Cryo EM data processing workflow using CryoSPARC**[2] **(supporting Figure 4). a)** Micrographs were subjected to motion correction, CTF correction, and template picking, yielding 985,743 particles that underwent 2D classification. These particles were sorted into two *ab initio* classes, separating a low-resolution (67,749 particles) from a well-resolved class (107,299 particles), which was further refined by 2D classification. Particles from both branches (175,048 and 93,495, respectively) were combined and processed through *ab initio* reconstruction into a single class, followed by box size adjustment and additional 2D classification, yielding a final particle stack of 78,699. This stack was used for non-uniform (NU) refinement under C6 and C1 symmetry, producing final reconstructions at 3.9 Å (C6) and 5.3 Å (C1) resolution. **b**) Gold-standard Fourier shell correlation (GSFSC) curves for the C6 map and C1 map.

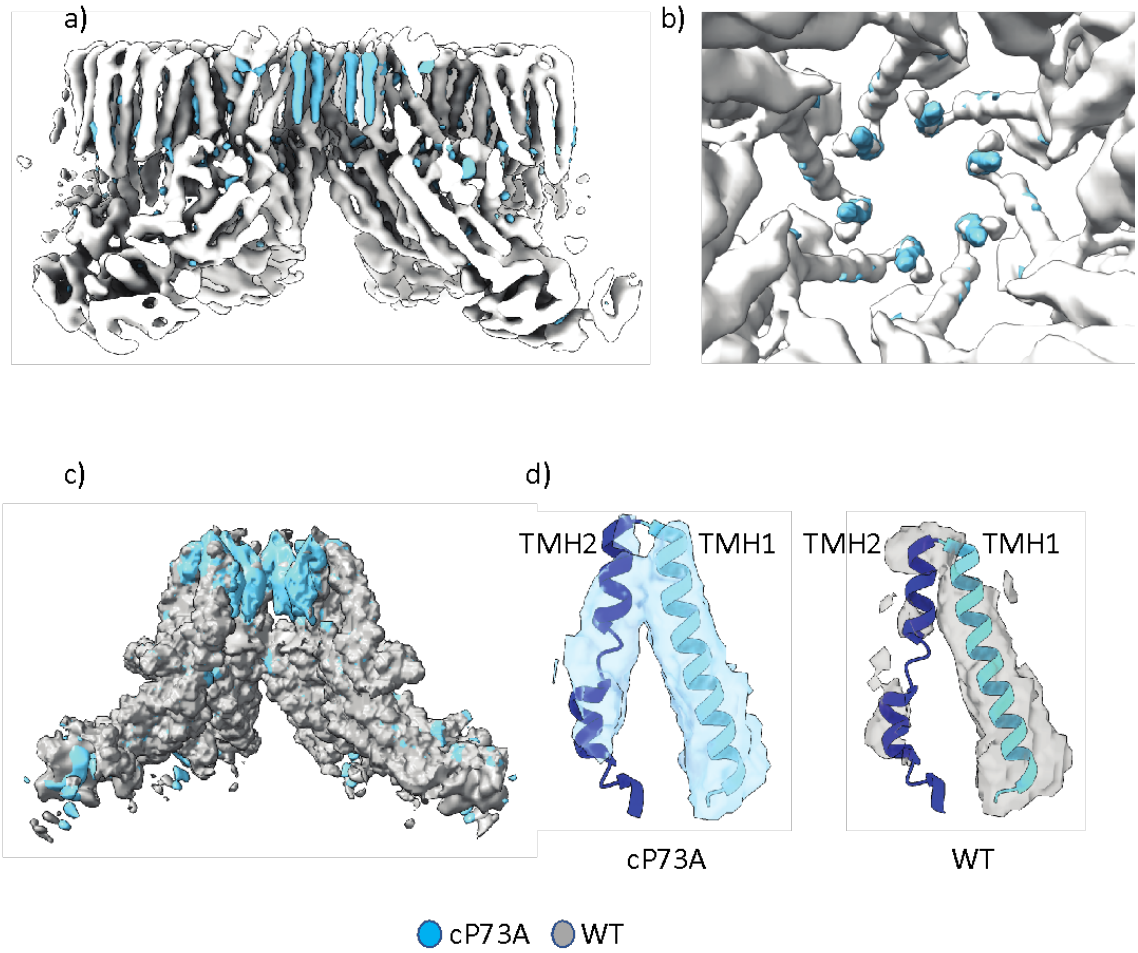


**Supplementary Figure 3. Changes in the ESX-5 pore architecture and dynamics induced by the cP73A mutation (supporting Figure 4).** Filtered difference map accounting for $EccC_5$ of ESX-5 (cP73A, blue) versus ESX-5 (WT, grey) a) in side view and b) zoomed top view (indicated by a dashed box in panel a). c) Overlay of cross-correlation maps calculated for half-maps, accounting for $EccC_5$ of ESX-5 (cP73A, blue) versus ESX-5 (WT, grey). d) cP73A TM section model overlaid with cP73A ESX-5 versus WT ESX-5 maps at cross-correlation map threshold levels that correspond to similar model-coverage.
